# Inflammatory profiles of transdiagnostic symptom dimensions in healthy females

**DOI:** 10.1101/2025.11.13.688353

**Authors:** Alicia J. Smith, Stacey Kigar, Quentin Dercon, Mary-Ellen Lynall, Konstantinos Ioannidis, Muzaffer Kaser, Caitlin Hitchcock, Tim Dalgleish, Camilla L. Nord

**Author notes:** Please address correspondence to:* Alicia Smith MRC Cognition and Brain Sciences Unit, University of Cambridge 15 Chaucer Road, Cambridge, CB2 7EF United Kingdom. Joint first authors (Alicia J. Smith, Stacey Kigar).

## Abstract

**Background:** Psychiatric disorders are increasingly conceptualised as heterogeneous categories with transdiagnostic underlying mechanisms that cut across multiple diagnoses and vary within a single diagnosis. Inflammation induced by psychosocial stress is one particularly potent example: previous research suggests that inflammatory profiles may correspond to symptom subgroups rather than traditional diagnostic categories. However, robust identification of transdiagnostic symptoms linked to specific inflammatory profiles remains rare. In this study, we examined the relationship between inflammatory profiles (at baseline and after a stress induction) and transdiagnostic symptom dimensions in females, who show higher prevalence of stress-related disorders such as anxiety and depression.

**Methods:** A modest but relatively homogenous healthy female sample, between the ages of 18 and 35, was recruited (N=26). We obtained venous blood samples, at baseline and after a combined physiological and social stress induction, to measure full blood counts, plasma cytokines and peripheral blood mononuclear cells (PBMCs; for cell stimulation and intracellular flow cytometry analysis). Participants completed a battery of psychiatric self-report questions, from which we modelled three transdiagnostic factor scores. Finally, we used Bayesian regressions to evaluate the predictive contribution of transdiagnostic factors to inflammatory markers (at baseline and stress-induced), as well as to the principal components of inflammatory measures (derived from a principal component analysis (PCA) of the inflammatory data).

**Results:** We identified specific relationships between inflammatory profiles and transdiagnostic symptom dimensions. Higher scores on a ‘social withdrawal’ factor were associated with greater baseline neutrophil (95% highest density interval (HDI) = [0.06, 1.32]; BF_10_ = 2.92) and monocyte counts (95% HDI = [0.29, 1.46]; BF_10_ = 19.08), whereas higher ‘anxious-depression’ scores were associated with a lower baseline monocyte count (95% HDI = [−0.96, −0.02]; BF_10_ = 1.81) and a greater inflammatory response to stress (e.g., change in neutrophil scores from pre- to post stress induction: (95% HDI = [0.18, 2.14]); BF_10_ = 5.66). We also found evidence for an association between the ‘social withdrawal’ factor and an immune principal component most strongly weighted by monocytes, basophils, and IL-6 (95% HDI = [−1.14, −0.01]; BF_10_ = 1.88).

**Conclusions:** We find preliminary evidence that different transdiagnostic psychiatric symptom dimensions map onto specific inflammatory profiles, both at baseline and after a stress induction. This represents a proof-of-principle for the use of data-driven and hypothesis-driven approaches to identify and link transdiagnostic factors with inflammatory changes, which may be of use to future studies with larger, clinical populations, or for testing in the context of stratified interventions.

## 1. Introduction

For over a century, psychiatric disorders have been organised into discrete categories based on formal taxonomic systems. Yet individuals with the same diagnosis often display significant heterogeneity in their symptoms and frequently present with comorbid conditions. Moreover, substantial overlap in symptomatology is observed across diagnostic categories (Dalgleish et al., 2020). Depression, for instance, is treated as a unitary disorder, but lacks a homogenous set of cardinal symptoms: in one study of 3,703 depressed outpatients meeting DSM-5 criteria, 1,030 unique symptom profiles were observed (Fried and Nesse, 2015), according to the current Diagnostic and Statistical Manual of Mental Disorders (DSM-5; American Psychiatric Association, 2013). Perhaps in part due to our current diagnostic taxonomy, most common psychiatric disorders appear aetiologically complex, with no (known) single process driving symptoms— whether neurobiological, psychological or environmental—a challenge that initiatives such as the NIMH Research Domain Criteria (RDoC; Insel et al., 2010) have sought to address by focusing on cross-diagnostic psychiatric domains.

Despite the longstanding importance of diagnoses in the assessment and treatment of mental ill-health, there is increasingly-mainstream support for research to move towards an alternative “transdiagnostic” approach that cuts across traditional boundaries, which might provide novel insights into the basis of psychiatric difficulties (Dalgleish et al., 2020; Pujji and Dinzeo, 2024). One of the most influential approaches, which emerges from the subfield of computational psychiatry, is the use of data-driven dimensionality reduction approaches in analyses of self-report questionnaires, and subsequent associations with neurocognitive mechanisms obtained via computational modelling (‘computational factor modelling’) (Wise et al., 2023); those are considered to have applications in classification, treatment selection and prediction of treatment response (see Huys et al., 2016). This approach was first applied by Gillan et al. (2016), who used an exploratory factor analysis on data from 9 commonly used psychiatric questionnaires to identify latent variables and derive transdiagnostic symptom scores for each participant. The three-factor structure identified—consisting of anxious-depression, compulsivity and intrusive thought, and social withdrawal—has since been replicated in several independent samples, supporting its robustness across populations and settings (Fox et al., 2025, 2023; Rouault et al., 2018; Seow and Gillan, 2020). This has facilitated the fast, robust identification of transdiagnostic symptom dimensions, even with abridged questionnaires (Wise and Dolan, 2020). In the past, these approaches have been confined to associations with cognitive measures but may hold broader potential in identifying transdiagnostic phenotypes associated with key biological mechanisms in psychiatry.

Both psychosocial stress and inflammation are implicated in psychiatric disorders, yet their effects cut across diagnostic boundaries, highlighting the inadequacy of current diagnostic nosology. Chronic psychosocial stress is a non-specific, but potent risk factor for a variety of mental health disorders (Brady and Sinha, 2005; Dinan, 2005). Both acute and chronic stress can induce peripheral inflammatory changes in humans (Fauci and Dale, 1974; Heidt et al., 2014; Kerr, 1956; Maydych et al., 2017) and in preclinical animal models (Curtin et al., 2009; Dhabhar et al., 1994; Heidt et al., 2014; Kigar et al., 2025; Lynall et al., 2020; Panzenhagen and Speirs, 1953). Inflammation, in turn, lacks specificity to any single diagnosis. For example, elevated plasma levels of interleukin-6 (IL-6)—a ‘keystone’ cytokine involved with both pro- and anti-inflammatory signalling (Hunter and Jones, 2015)—are consistently reported in people with depression (Khandaker et al., 2018; Osimo et al., 2020), but are also present in eating disorders (Dalton et al., 2018), psychosis, and post-traumatic stress disorder (PTSD) (Lindqvist et al., 2014; Passos et al., 2015). We recently confirmed case-control differences in IL-6, as well as other peripheral cytokines and immune cell types, in depressed individuals (Lynall et al., 2020). Moreover, in this study, we found 4 depression subtypes delineated by unique inflammatory profiles, which corresponded to symptom severity. Broadly speaking, immune dysregulation has been reported in psychiatric disorders sharing symptoms with depression, including anxiety disorders, bipolar disorder, schizophrenia, and PTSD (Costello et al., 2019; Goldsmith et al., 2023, 2016; Munkholm et al., 2013), and genetic risk analyses implicate adaptive immune cells in the pathogenesis of multiple psychiatric disorders, including depression (Lynall et al., 2022).

Decades of research in animals have described bidirectional communication between the brain and immune system, influencing behaviour. For example, the brain exhibits top-down regulation of the immune system (Schiller et al., 2021). One well-described pathway for this is via activation of the hypothalamic-pituitary-adrenal (HPA) axis; HPA activation leads to the release of glucocorticoids, which have potent but mixed effects on both immune cells and cytokines (Herkenham and Kigar, 2017; Segerstrom and Miller, 2004). Conversely, experimentally-induced inflammation drives behavioural alterations, including social withdrawal and diminished motivation to engage or react to incentives (Blank et al., 2016; Dantzer and Kelley, 2007). These behavioural changes, collectively termed ‘sickness behaviours’, share some symptom overlap with depression (Dantzer and Kelley, 2007; Eisenberger et al., 2017; Slavich, 2020). In human and laboratory animals, acute administration of a low-dose endotoxin potently induces cytokines transiently and increases social anhedonia and anxiety (Eisenberger et al., 2010; Lasselin et al., 2016). Accordingly, a recent study revealed that specific transdiagnostic symptoms analogous to sickness behaviours are responsive to anti-inflammatory drugs (Lee et al., 2020).

Thus, immune dysregulation may be best understood as a transdiagnostic mechanism in psychiatry (Franklyn et al., 2022; Majd et al., 2020). Novel approaches drawing on this transdiagnostic understanding are needed to advance mechanistic understanding and enable more personalised care. While some immunopsychiatry studies have focused on immune cells (Lynall et al., 2020; Maes, 2011; Segerstrom and Miller, 2004), many psychiatry studies examine plasma cytokine levels, e.g., in relation to case-control differences (Chamberlain et al., 2019; Lynall et al., 2020; Miller et al., 2014), symptom severity (Daria et al., 2020; Keeler et al., 2021; Yeon et al., 2017), or longitudinal risk (Khandaker et al., 2014; Lee et al., 2020; van Eeden et al., 2021; Van Eeden et al., 2020). Collectively, this work has yielded important insights into putative mechanisms associated with mental illness. However, deriving inferences from basal cytokine levels can be challenging given high variation between individuals and that circulating concentrations may be well below the limit of detection for most commercial assays (Wu et al., 2017). In addition, basal inflammatory markers appear to be sensitive to lifestyle and health factors such as smoking and body mass index (BMI) (Vogelzangs et al., 2016). Investigation of cytokine output in response to an immune challenge such as lipopolysaccharide (LPS) stimulation circumvents some of these issues, appears to reflect heritable differences in immune system function, and may correlate more strongly with symptom severity (De Craen et al., 2005; van der Linden et al., 1998; van Eeden et al., 2021; Van Eeden et al., 2020; Vogelzangs et al., 2016). This suggests LPS stimulation of white blood cells may unmask additional mechanisms relevant to psychiatry, complimenting work on basal inflammatory markers.

The purpose of this study was to investigate the relationship between transdiagnostic symptom dimensions identified in previous work (Gillan et al., 2016; Wise and Dolan, 2020) and several components of inflammation in healthy adult females, measured at baseline and following acute stress. In this proof of concept study, we chose to focus on females for two reasons: first, to maintain a relatively homogeneous sample population due to reported sex differences in both the laboratory stress response (Henze et al., 2021; Quaedflieg et al., 2013) and in psychiatric factor analysis (Saunders et al., 2023); secondly, because women are disproportionately affected by stress-related psychiatric disorders like depression and anxiety (Farhane-Medina et al., 2022; Kigar and Auger, 2013). Consistent with previous studies employing the same transdiagnostic methods, in this study we capture inflammatory relationships with the variation of symptoms present in a non-clinical sample. Although this design precludes direct conclusions about clinical populations, it provides an opportunity to explore underlying biological associations in a controlled, proof of concept manner, without confounds such as psychiatric medication. Here, we report stress-related changes in *ex vivo* peripheral white blood cell counts and plasma cytokine levels, as well as in vitro intracellular cytokine levels following LPS stimulation, to better understand how inflammatory responses relate to transdiagnostic dimensions of mental health.

## 2. Methods

### 2.1 Participants

To mitigate against known immunological differences in the population (Klein and Flanagan, 2016), we restricted recruitment according to the following criteria, (1) female sex assigned at birth, (2) age 18-35 years old (mean age = 26 ± 4.7), (3) no major medical conditions (including but not limited to: cancer, diabetes, lupus, Huntington’s disease, etc.), (4) no current drug treatments that alter cognition, mood, or the immune system (including but not limited to non-steroidal anti-inflammatory medication, antidepressant, or antipsychotic medication, etc.), (5) willing to refrain from exercise for 72 hours and fast for 12 hours preceding the study. Other inclusion criteria were (6) normal or corrected-to-normal vision, and (7) fluent or native English speakers. All criteria were listed on the recruitment poster and confirmed during a screening call with the researcher (AJS) prior to enrolment. All subjects provided written informed consent prior to taking part. Participants were paid a fixed rate of £10 per hour of in-person testing, plus payment for travel. Ethics were approved by the University of Cambridge Human Biology Research Ethics Committee (HBREC.2020.40).

### 2.2 Experimental Procedure

After an initial telephone appointment to determine eligibility, participants were sent a link to complete cognitive tests from home within the week leading up to their in-person assessment at Cambridge Clinical Research Centre (CCRC) located at Addenbrookes Hospital. The at-home cognitive testing included questionnaires used to determine scores on three transdiagnostic dimensions (see Methods 2.2.1). Catch questions were asked at multiple time points to detect inattentive responses in line with best practice for online data collection.

On the day of their in-person assessment, participants provided informed consent on arrival, before fasting venous blood samples were taken. Next, participants undertook the Maastricht Acute Stress Test (MAST; Smeets et al., 2012) before completing cognitive testing, which comprised task-based assessments, the results of which have been reported elsewhere (Smith et al., 2024). Participants completed a subjective stress questionnaire immediately after the stress induction, and venous blood samples were collected 45 minutes thereafter, representing the earliest time point at which a rise in plasma IL-6 is thought to be detectable following acute laboratory stress (Endrighi et al., 2016). Post-stress blood sampling was done prior to peak IL-6 plasma release (approximately 75 minutes after the stressor) to maximise the detection of intracellular IL-6 staining. All samples were taken between 08:00 and 11:00 am. For each time point, a 3ml subset of blood was taken for hospital measurement of absolute blood cell counts. The remaining blood was used for immunophenotyping and biomarker collection. The experimental procedure is illustrated in **Figure 1**.

**Figure 1.**
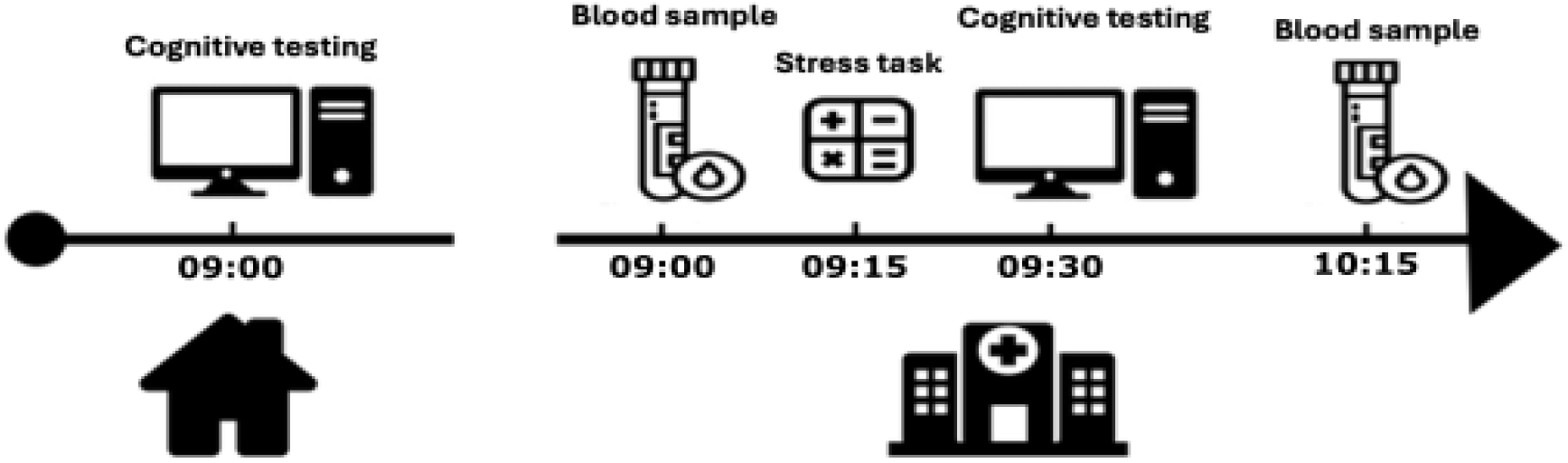
Experimental procedure with approximate timings. Participants completed a battery of mental health questionnaires at home prior to their in-person assessment at the Cambridge Clinical Research Centre. On the day of the in-person assessment, fasting venous blood samples were taken on arrival, after which participants completed the Maastricht Acute Stress Test. Forty-five minutes after the stress test, blood samples were taken for a second time.

#### 2.2.1 Questionnaires

Three transdiagnostic dimensions, namely ‘anxious-depression’, ‘social withdrawal’, and ‘compulsivity and intrusive thought’, have previously been identified using a factor analysis on a set of 209 questionnaire items in work by Gillan et al. (2016). These dimensions have widely been used to examine underlying neurocognitive mechanisms (Wise et al., 2023). We employed a previously-established method (Wise and Dolan, 2020) to identify a subset of questions that could be used to predict subjects’ transdiagnostic scores more efficiently (**Figure 2**). Specifically, we applied a multi-target Lasso regression to predict factor scores from a full set of question responses collected in a study by Rouault et al. (2018). Differing numbers of question coefficients were fixed to zero and five-fold cross-validation was then used to assess the predictive accuracy of these subsets, whereby the model was trained on 80% of the data and tested on the remaining 20%. Retaining a set of 78 questions represented a good compromise between the number of questions and predictive accuracy, with R^2^ values of ≥ 0.9 for all three factors. Consequently, participants in the present study completed 78 mental health question items, which were then used to estimate their individual scores on the three transdiagnostic dimensions.

**Figure 2.**
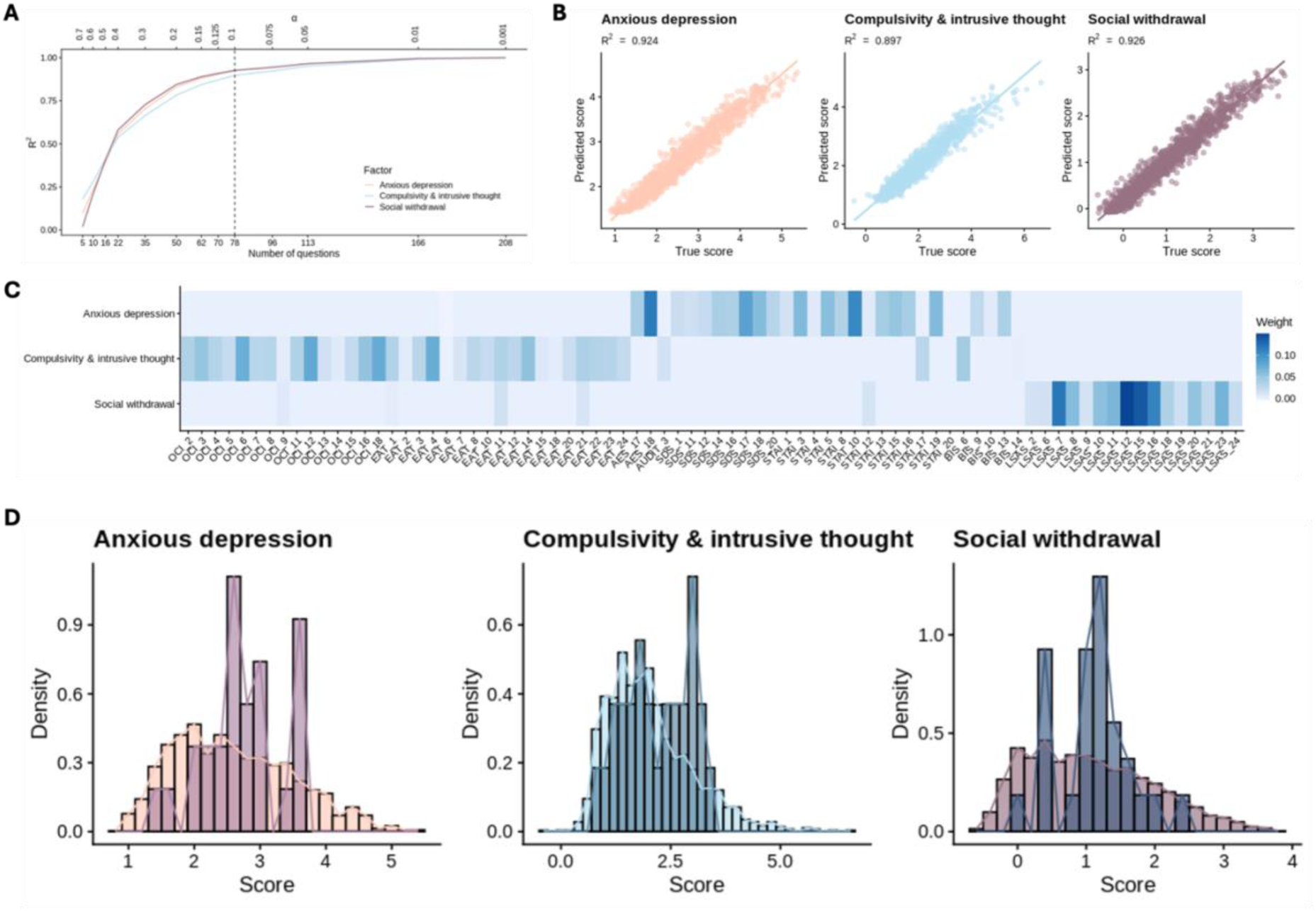
Results from the questionnaire reduction method for measuring three transdiagnostic factors. Using multi-target lasso regularised regression, factor scores were predicted from responses to subsets of questions in the original dataset obtained in the study by Gillan et al. In this model, the alpha value determined the degree of regularisation and predictive accuracy was assessed using five-fold cross-validation, with R^2^ averaged across the five folds. (**A**) Alpha and R^2^ values were plotted allowing us to select a point at which there was a good compromise between predictive accuracy and number of questions retained. An alpha value of 0.1 was selected for the present study, resulting in a subset of 78 questions out of the initial 209. (**B**) True factor scores produced using the full questionnaire dataset from the Gillan et al. study plotted against predicted factor scores produced by the selected model. The selected model showed good predictive accuracy (R^2^ ≥ 0.9) for all three factors. (**C**) Heatmap demonstrating the weights of each of the retained questions for each transdiagnostic factor. (**D**) Distributions of factor scores for the participants in the present study (darker bars) overlaid on distributions of factor scores obtained in the study by Gillan et al. study (lighter bars).

#### 2.2.2 Stress induction

The Maastricht Acute Stress Test (MAST) is a laboratory procedure that involves both a physical and psychosocial stressor, and has previously been shown to reliably elicit autonomic, glucocorticoid, and subjective psychological reports of stress responses (Smeets et al., 2012). The task, lasting 12 minutes, requires participants to alternate between submerging their hand in ice water and answering mental arithmetic questions in front of an assessor providing negative feedback (e.g., ‘go faster!’).

#### 2.2.3 Sample collection

Approximately 23ml of EDTA fasting venous blood was taken prior to- and 45 minutes after completing the stress induction. This represents the earliest timepoint with a detectable rise in plasma IL-6 following acute laboratory stress (Endrighi et al., 2016). Post-stress blood sampling was done prior to peak IL-6 plasma release (peak occurs approximately 75 minutes after the stressor) to maximize detection of intracellular IL-6 staining. To minimise confounding by diurnal variation in markers, all samples were taken between 08:00 and 11:00am. For each timepoint, a 3ml aliquot of blood was used for measurement of absolute blood cell counts (neutrophils, eosinophils, basophils, lymphocytes and monocytes) at a hospital clinical laboratory. The remaining blood was used for immunophenotyping and biomarker assays (see Methods 2.2.4 - 2.2.6).

#### 2.2.4 Plasma isolation and cytokine analysis

Immediately upon laboratory receipt, 10ml of EDTA blood was centrifuged for plasma aliquot collection. Samples were stored at −80°C until completion of the study. Once all participant samples were collected, aliquots were thawed for single batch analysis via the V-PLEX Proinflammatory Panel 1 multiplex assay (MSD, Catalogue#: K15049D-1) with a MesoScale Discovery Sector s600 plate reader. This permitted simultaneous assessment of the following cytokines: interferon-γ (IFN-γ), interleukin (IL)-1β, IL-2, IL-4, IL-6, IL-8, IL-10, IL-12p70, IL-13, and tumor necrosis factor-α (TNF-α). Intra-assay coefficient variations (CVs) were not calculated due to their only being two sample plates; inter-assay CVs are presented in **Supplemental Table S1**. Four of the analysed cytokines (IL-1β, IL-2, IL-4, and IL-13) were excluded from the analysis due to more than 20% of the values being below the lower limit of detection (LLOD) as deterioration of imputation method accuracy has been shown to occur for this level of data missingness (Liu and Brown, 2013). For cytokines with less than 20% of the values below the LLOD, maximum likelihood estimates (MLE) were used to impute values for left-censored data given that this method is shown to reduce bias compared to other replacement methods (Palarea-Albaladejo and Martín-Fernández, 2015), including the commonly used 0.5*LLOD method (Zhang and Guo, 2012).

#### 2.2.5 Peripheral blood mononuclear cell (PBMC) isolation and immune challenge

All remaining blood was gently layered onto sterile Ficoll (Cytiva, Catalogue#: 17144003) for PBMC isolation. The resulting PBMCs were then plated in duplicate wells for each condition (stimulated or non-stimulated for both baseline and post-stress conditions) at a density of 4×10^5^ cells per well in RPMI 1640 with L-glutamine (Sigma-Aldrich, Catalogue#: R8758-500ML) supplemented with 100 IU/ml penicillin and 100 mg/ml streptomycin (Sigma-Aldrich, Catalogue#: P4458-100ML). For each timepoint per participant, an additional well was generated for flow cytometry staining controls (see Methods 2.2.6). Each sample well was treated with GolgiSTOP (BD, Catalogue#: 51-2092KZ) to trap cytokines intracellularly. Stimulated wells were treated with 50ng of lipopolysaccharide (LPS), which mimics the cell wall of gram-negative bacteria (Sigma-Aldrich, Catalogue#: L-2880-10MG) and elicits an immune response. Non-stimulated wells received an equivalent volume of vehicle. After gently mixing on a plate shaker, the cells were incubated at 37°C for 4 hours. At the end of the incubation, cells were transferred to 5ml polystyrene tubes and washed to prepare for flow cytometry staining.

#### 2.2.6 Intracellular IL-6 staining and flow cytometry

Each sample was stained for extracellular markers using the antibody cocktail shown in **Supplemental Table S2** (20 minutes, 4°C). Cells were then washed and fixed for 30 minutes at room temperature using Fixation/Permeabilization solution (FoxP3/TF kit, Thermo Fisher, Catalogue#: 00-5523-00). After fixation, cells were diluted with the kit’s Wash Buffer and left overnight at 4°C. The next day, cells were washed and resuspended in residual buffer. IL-6-PE (BD, Catalogue#: 559331) was added to each sample, except for the control samples mentioned in section 1.2.5; these became fluorescence minus one (FMO) controls used to define positive and negative intracellular IL-6 staining for each individual participant. The intracellular staining reaction proceeded at room temperature for 20 minutes. Cells were then washed before data acquisition on a BD Symphony flow cytometer. After data acquisition, manual flow cytometry gating was performed using FlowJo software (BD). 6 unique cell populations (monocytes, B cells, T cells, NK cells, NKT cells, and dendritic cells) were identified according to the gating strategy shown in **Supplemental Figure S1**. Each cell type was assessed for intracellular IL-6 staining; only monocytes and B cells showed any positive staining, and thus the results from the other cell types are not shown. Positive IL-6 staining was calculated as a percentage relative to parent cell type. Duplicates for each time point and condition were averaged, then the stimulated values were normalized to the non-stimulated values for presentation as fold-change at each time point (baseline and post-stress).

### 2.3 Statistical analyses

Multiple univariate analyses were used to examine baseline to post-stress differences in peripheral blood cell counts, cytokine concentrations, and intracellular IL-6 staining. Bayesian regressions assessed evidence for relationships between inflammatory measures and transdiagnostic symptom dimensions, allowing us to quantify uncertainty and incorporate prior knowledge, which is particularly advantageous given the small sample size in this study. We report 95% Highest Density Intervals (HDIs), which represent the range within which the parameter values fall with 95% probability. A 95% HDI that does not include zero is interpreted as evidence for a non-zero effect, indicating that zero is not among the most credible values for the parameter given the data.

All three transdiagnostic factors were entered as independent variables into the regression model, predicting each inflammatory cytokine or cell count at baseline, as well as the change from baseline to post-stress levels (from here on, referred to as *stress-induced inflammation*). MinMax scaling was used to normalise the dependent parameters prior to model fitting. Models were built using Bambi (Capretto et al., 2022) and fit using Markov chain Monte Carlo sampling, with 2,000 warm-up draws and 8,000 sampling draws per chain.

Multiple comparisons corrections using the Bonferroni procedure were performed for univariate analyses and correlations to control the false discovery rate; no multiple comparison corrections were used for regression modelling due to this being unnecessary and incompatible with the Bayesian approach used in this study (Gelman et al., 2012).

A principal component analysis (PCA) was conducted to explore underlying dimensions of covariation across the inflammatory variables. All variables were centred and scaled prior to analysis, such that the mean was equal to zero and the standard deviation was equal to one for each variable. Given the exploratory nature of this analysis and the relatively small sample size, the PCA was used cautiously as a data reduction method. Sampling adequacy was evaluated using the Kaiser-Meyer-Olkin (KMO) measure and Bartlett’s Test of Sphericity to assess the appropriateness of the PCA. The overall KMO value was 0.50, indicating a minimum acceptable level of sampling adequacy for a PCA. Barlett’s Test was significant (*p* < 0.05), suggesting that the variables were sufficiently correlated for PCA. The number of principal components to retain was selected based on the cumulative variance explained, with the first three components accounting for approximately 70% of the total variance.

Statistical analyses were performed using RStudio and Python. Code is available at (https://github.com/aliciasmith1/transdiagnostic-inflammatory-profiles).

## 3. Results

### 3.1 Descriptive characteristics

Baseline and post-stress absolute blood cell counts of 5 white blood cell types were available for a sample of 23 participants, baseline and post-stress concentrations of 6 inflammatory plasma cytokines were available for 26 participants, and baseline and post-stress intracellular IL-6 staining for 2 cell types were available for 16 participants. Each analysis was conducted using the maximum available data. Sample characteristics are presented in **Table 1**.

**Table 1.** Sample characteristics.

|  | **Total**<br>**(N=26)** |
| --- | --- |
| **Gender, number (%)** |  |
| Male | 0 (0 %) |
| Female | 25 (96 %) |
| Non-binary | 1 (4 %) |
| **Age (years)** |  |
| Mean (SD) | 25 (4.6) |
| **Ethnicity, number (%)** |  |
| White | 22 (85 %) |
| Asian | 2 (8 %) |
| Mixed | 2 (8 %) |
| Other | 0 (%) |
| **Subjective socioeconomic status (/9)** |  |
| Mean (SD) | 5.7 (1.0) |

### 3.2 Baseline to post-stress differences in peripheral white blood cell counts

We first conducted univariate (frequentist) regression analyses and non-parametric group comparisons to examine changes in peripheral white blood cell counts from baseline to post-stress (**Figure 3A**), applying a Bonferroni-corrected significance threshold of *p* < 0.01 to account for 5 comparisons. We observed stress-associated increases in neutrophil absolute counts (t(21) = 4.95, \*\*\**p* < 0.001, d = 1.06, 95% CI [−0.77, −0.32]), as well as stress-associated decreases in lymphocyte counts (t(21) = −3.91, \*\**p* = 0.001, d = 0.83, 95% CI [0.14, 0.46]) and eosinophil counts (Wilcoxon Signed-Ranks: Z = −3.16, \*\**p* = 0.002, d = 0.81, 95% CI [0.01, 0.04]). No significant changes were observed in monocyte (t(21) = 0.30, *p* = 0.77, d = 0.06, 95% CI [-0.02, 0.03]) or basophil counts (Wilcoxon Signed-Ranks: Z = 0.17, *p* = 0.89, d = 0.05, 95% CI [-0.01, 0.01]).

**Figure 3.**
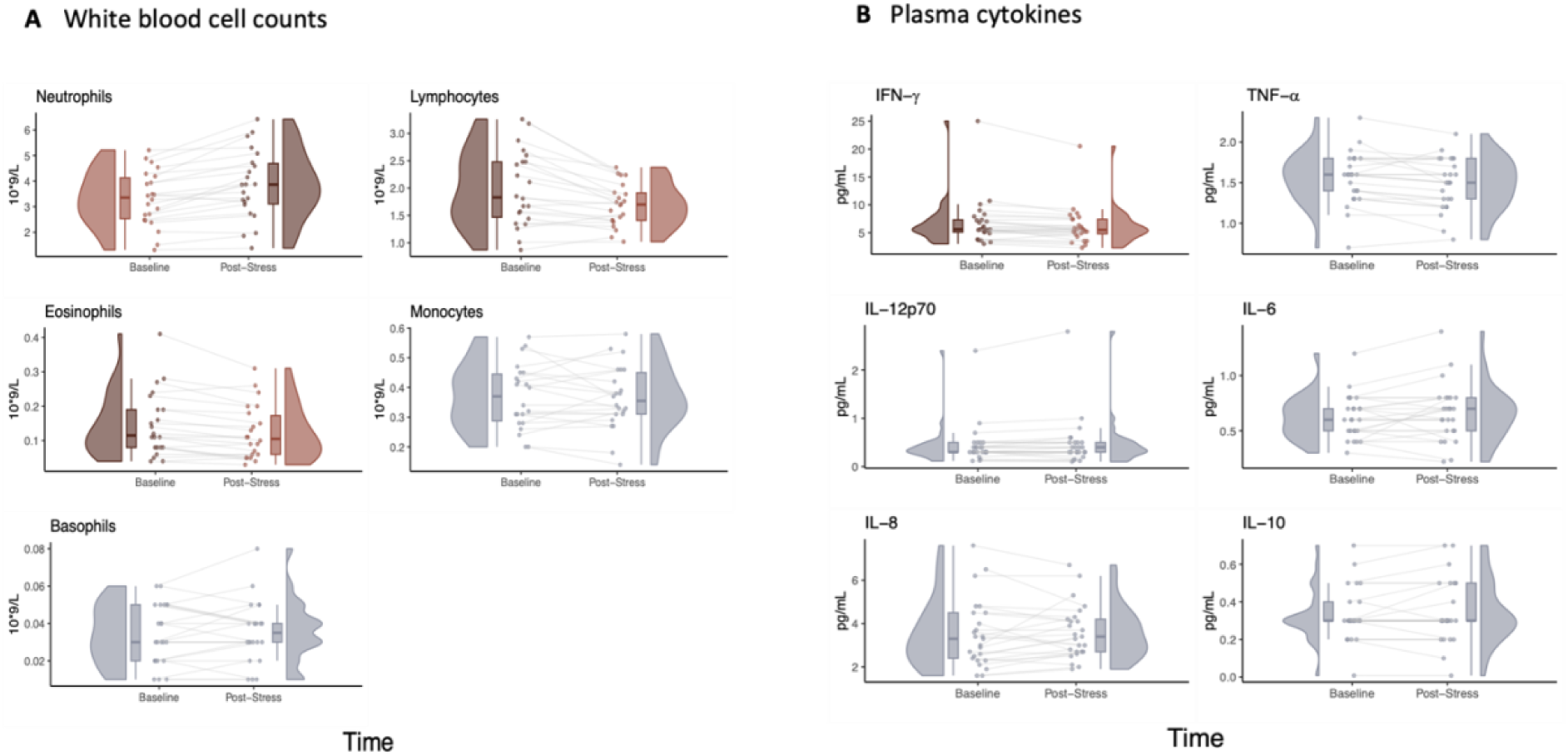
Baseline to post-stress changes in blood cell counts, plasma cytokine concentrations. Plots displayed in red indicate that there was a significant difference (FDR *p* < .05) in (**A**) white blood cell counts or (**B**) plasma cytokines between baseline and post-stress, with dark red plots indicating the time point at which the measurement was higher and lighter red plots indicating the time point at which the measurement was lower. Scatter points show individual subjects’ measurements at baseline and post-stress, kernel density plots illustrate the distribution of the values and box plots depict the median and interquartile range.

### 3.3 Baseline to post-stress differences in plasma and intracellular cytokines

We then examined changes in plasma cytokine levels from baseline to post-stress (**Figure 3B**), applying a Bonferroni-corrected significance threshold of *p* < 0.0083 to account for 6 comparisons. We observed a significant decrease in IFN-γ levels following stress (t(24) = - 3.01, *p* = *0.006, d = 0.60, 95% CI [0.25, 0.95]).

No significant changes were observed for IL-12p70 (Wilcoxon Signed-Ranks: Z = 1.57, *p* = 0.12, d = 0.50, 95% CI [-0.15, 0.01]), IL-6 (t(24) = 1.30, *p* = 0.21, d = 0.26, 95% CI [-0.10, 0.02]), IL-8 (t(24) = 0.46, *p* = 0.65, d = 0.09, 95% CI [-0.42, 0.27]), IL-10 (Wilcoxon Signed-Ranks: Z = 0.53, *p* = 0.63, d = 0.20, 95% CI [-0.10, 0.10]) or TNF-α (Wilcoxon Signed-Ranks: Z = −2.33, *p* = 0.02, d = 0.63, 95% CI [0.05, 0.15]).

Plasma IL-6 reflects the combined output of IL-6 from multiple immune and other cell types. To investigate cell-specific effects of stress on IL-6 production, we tested for intracellular IL-6 in monocytes and B cells. We observed a significant increase in monocyte intracellular IL-6 following stress (Wilcoxon Signed-Ranks: Z = 2.44, \**p* = 0.012, d = 0.72, 95% CI [−15.64, - 2.23]) (**Figure 4**), but no stress-associated increases in B cell IL-6 (t(14) = 1.90, *p* = 0.08, d = 0.49, 95% CI [-0.66, 0.04]).

**Figure 4.**
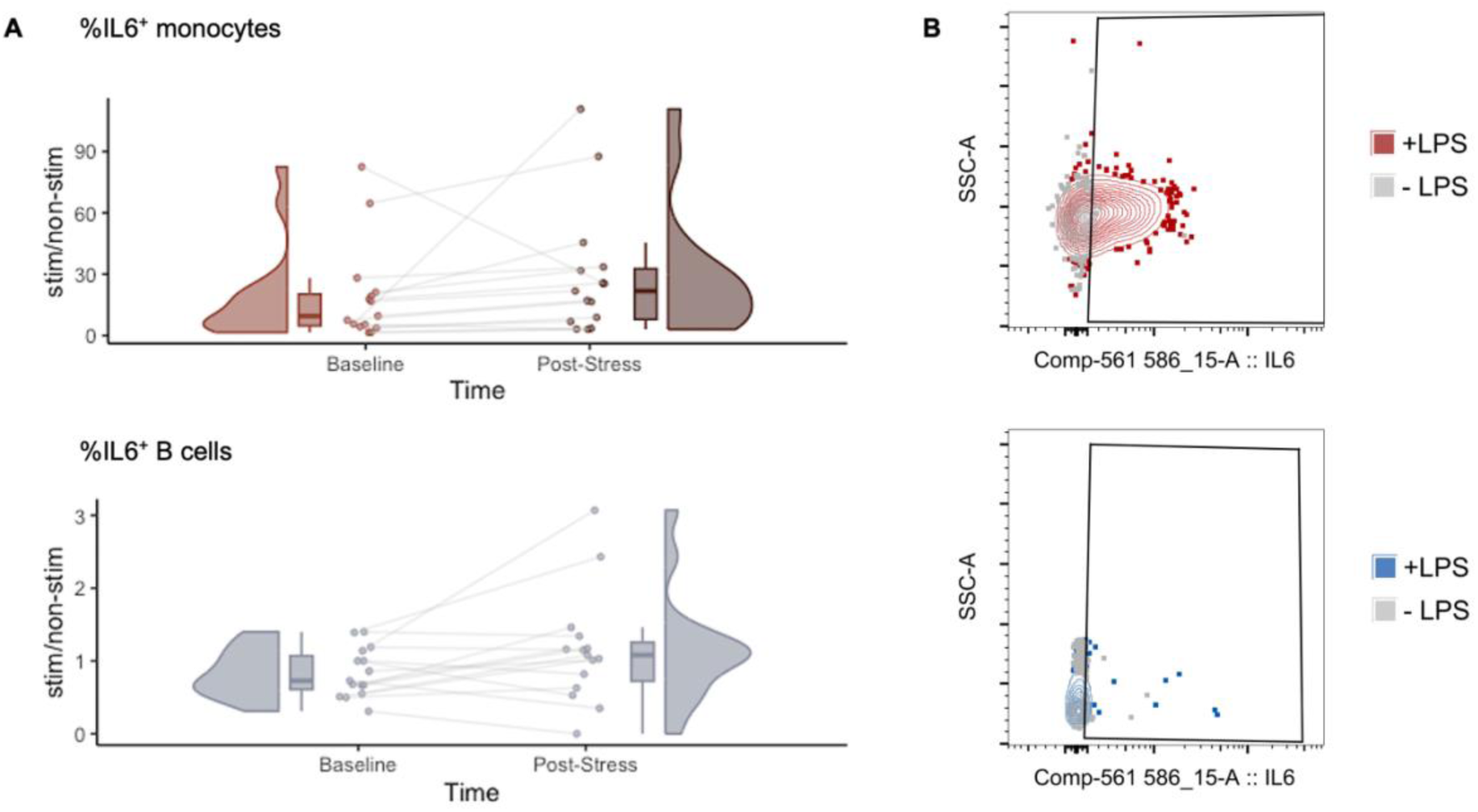
Effects of laboratory stressor on cell specific IL-6 production. (**A**) At 45 minutes post-laboratory stress, stimulated monocytes produce more IL-6 compared to baseline; stimulated B cells do not show a significant change in IL-6 production. Plots displayed in red indicate a significant difference (FDR *p* < .05) in intracellular IL-6 staining between baseline and post-stress, with dark red indicating the time point at which the measurement was higher and lighter red indicating the time point at which the measurement was lower. Scatter points show individual subject’s measurements at baseline and post-stress, kernel density plots illustrate the distribution of the values and box plots depict the median and interquartile range. Values represent fold-change enrichment of intracellular IL-6 staining (calculated as percent positive relative to the parent cell population) between lipopolysaccharide (LPS)-stimulated and non-stimulated cells for each participant sample. (**B**) Representative bivariate plots corresponding to the cell type shown at left showing flow cytometric data for LPS stimulated (+LPS) and vehicle treated (-LPS) cells. Colour reflects level of significance in (**A**). A fluorescence minus one (FMO) control sample, which lacked IL-6 antibody staining, was used to define the true negative population for each participant sample. The x-axis shows intracellular IL-6 staining, whereas the y-axis shows side scatter properties (indicative of intracellular complexity).

### 3.4 Transdiagnostic dimensions & inflammation at baseline and in response to stress

Using Bayesian regressions, we examined whether each transdiagnostic factor (‘social withdrawal’, ‘anxious-depression’, ‘compulsivity and intrusive thought’) was associated with baseline measures of white blood cell counts, plasma cytokines, and intracellular IL-6 staining (**Figure 5**). We found strong evidence for a positive association between ‘social withdrawal’ and baseline monocytes (95% HDI = [0.29, 1.46]; BF_10_ = 19.08), and anecdotal evidence for a positive association between ‘social withdrawal’ and baseline neutrophils (95% highest density interval (HDI) = [0.06, 1.32]; BF_10_ = 2.92), with HDIs that did not include zero. We additionally found anecdotal evidence for a negative association between ‘anxious-depression’ and baseline monocyte counts (95% HDI = [-0.96, −0.02]; BF_10_ = 1.81), as well as between ‘compulsivity and intrusive thought’ and plasma IL-12p70 (95% HDI = [−0.69, −0.02]; BF_10_ = 1.39). All other associations were weak, with the 95% HDIs overlapping with zero.

**Figure 5.**
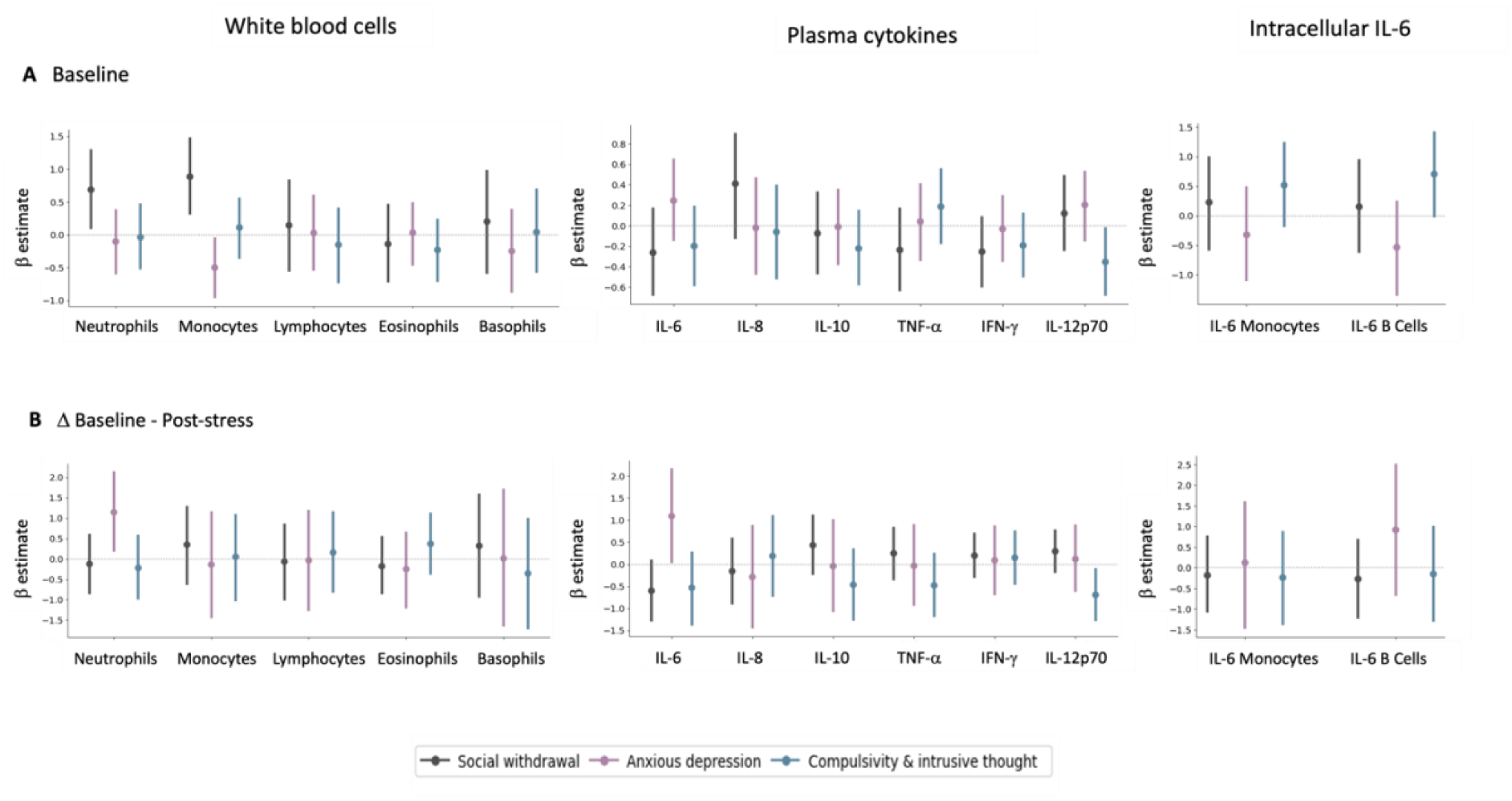
Results from Bayesian regression models relating three transdiagnostic dimensions to (A) baseline immunological measures, and (B) immunological responses to stress, in terms of white blood cell counts, cytokine levels, and intracellular IL-6 staining. Points illustrate the mean of the posterior distribution, along with error bars representing the 95% highest density interval of the regression coefficient for each predictor.

We also examined the extent to which the three transdiagnostic factors were associated with stress-induced inflammation, measured as the absolute difference in white blood cell counts, plasma cytokine concentrations, and intracellular IL-6 staining between baseline and post-stress (**Figure 4**). We found moderate evidence for a positive association between ‘anxious-depression’ and both post-stress neutrophil count change (95% HDI = [0.18, 2.14]; BF_10_ = 5.66) and post-stress plasma IL-6 level change (95% HDI = [0.03, 2.17]; BF_10_ = 3.59). We additionally found moderate evidence for a negative association between ‘compulsivity and intrusive thought’ and post-stress plasma IL-12p70 change (95% HDI = [−1.29, −0.09]; BF_10_ = 3.49). No other associations provided evidence of directional effects, with the 95% HDIs overlapping with zero. Regression coefficients are reported in **Supplemental Tables S3** and **S4**, including the mean of the posterior distribution and the 95% HDIs.

### 3.5 Correlations and PCA of inflammatory markers

Next, we analysed correlations between transdiagnostic variables and baseline inflammation, as well as with stress-induced inflammatory levels (calculated as the absolute difference in white blood cell counts, cytokines, or intracellular IL-6 staining between baseline and post-stress) (**Figure 6**). Analyses were Bonferroni-corrected to a significance threshold of *p* < 0.0004. We found strong evidence for a positive correlation between baseline levels of lymphocytes and basophils (rho = 0.70, *p* = 0.0003). Notably, plasma IL-6 and monocyte IL-6 levels were non-correlated at baseline (rho = −0.09, *p* = 0.74), but the stress-induced change in plasma and monocyte IL-6 concentrations showed a trend to correlation (rho = −0.64, *p* = 0.01).

**Figure 6.**
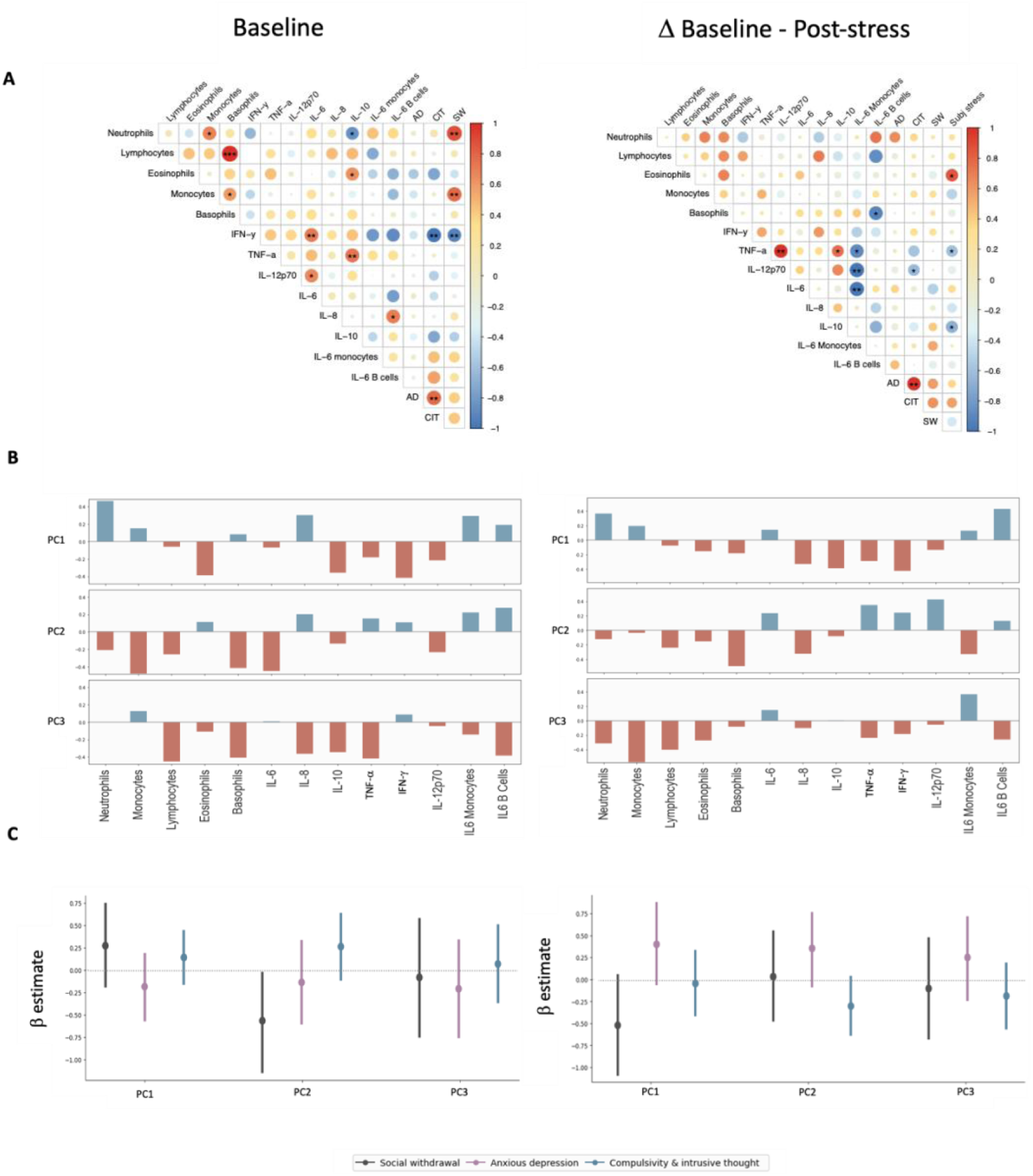
(A) Correlogram displaying Spearman’s correlations between the three transdiagnostic dimensions and immunological variables measured at baseline (left panel), and immunological variables in terms of their baseline to post-stress differences (right panel). Size and colour of circles signify the degree and directionality of the correlation coefficient (Spearman’s ρ). The correlogram is ordered using the First Principal Component (FPC) order argument, indicating the features that explain the maximum variance in the model. Asterisks indicate uncorrected significance levels (* *p* < 0.05, ** *p* < 0.01, *** *p* < 0.001); however, only a baseline correlation between lymphocytes and monocytes met the Bonferroni-corrected threshold for statistical significance (*p* < 0.0004). **(B) Bar diagram displaying variable loadings from a principal components analysis (PCA) of cellular, plasma and intracellular IL6 inflammatory variables.** Individual bar diagrams show loadings of each inflammatory variable measured at baseline (left panel), and in terms of their pre- to post-stress change (right panel), on immune principal component (PC) 1, PC2 and PC3. High absolute values indicate that the variable has a strong influence on the component. Values close to 0 indicate that their influence on the component is weak. A positive loading (blue bar) indicates that the variable and principal component are positively correlated. A negative loading (red bar) indicates that they are negatively correlated. **(C) Results from Bayesian regressions relating three transdiagnostic dimensions to the three principal components derived from measurements of inflammatory variables measured at baseline (left panel), and in terms of their pre- to post-stress change (right panel).** AD, Anxious-depression; SW, Social withdrawal; CIT, Compulsivity and intrusive thought; IFN-γ, Interferon-γ; TNF-α, Tumor Necrosis Factor-α; IL-6, Interleukin-6; IL-8, Interleukin-8; IL-10, Interleukin-10.

A principal components analysis (PCA) was conducted to explore patterns of covariation among the inflammatory variables (**Figure 6**, see Methods). Based on the explained variance, 3 principal components were extracted (**Supplemental Figure S2**). Together, they accounted for 70% of the total variance. The first immune principal component (PC1) was most strongly weighted by neutrophils, eosinophils, IL-8, IL-10 and IFN-γ, and explained 30% of the total variance. The second component was most strongly weighted by monocytes, basophils and IL-6 and accounted for 23% of the total variance. The third component was most strongly weighted by lymphocytes, basophils, IL-8, IL-10, TNF-α and IL-6^+^ B cells, and explained 18% of the total variance. **Figure 6B** shows item loadings on each principal component, where a high loading value (close to 1 or −1) implies that the variable strongly influences the component, and the sign of the component (either + or -) indicates whether a variable is positively or negatively correlated with the principal component. Using Bayesian regressions, we examined whether each transdiagnostic factor (‘social withdrawal’, ‘anxious-depression’, ‘compulsivity and intrusive thought’) was associated with baseline measures of immune PC1, PC2 and PC3. We found anecdotal evidence for a negative association between the ‘social withdrawal’ factor and immune PC2 (95% HDI = [−1.14, −0.01]; BF_10_ = 1.88). No other relationships strongly supported directional effects, with the 95% HDIs overlapping zero (**Supplemental Table S5**).

We additionally ran a PCA summarising the stress-induced inflammatory variables (**Figure 6**). We retained the first 3 principal components based on the component variance (**Supplemental Figure S2**), which collectively explained 59% of the total variance-covariance between the inflammatory variables. **Figure 6B** displays the loadings for each variable on the first 3 principal components. Immune PC1 accounted for 22% of the total variance-covariance and was most strongly weighted by neutrophils, IL-8, IFN-γ and IL-6^+^ B cells. Immune PC2 also accounted for 22% of the total variance-covariance and was most heavily weighted by basophils, IL-8, TNF-α, IL-12p70 and IL-6^+^ monocytes. Lastly, immune PC3 accounted for 15% of the total variance-covariance and was most strongly weighted on monocytes, lymphocytes and IL-6^+^ monocytes. Bayesian regressions were used to explore relationships between the three transdiagnostic factors and baseline-to-post-stress changes in the three principal components. We did not find strong evidence for any relationships between these variables, with all 95% HDIs overlapping with zero (**Supplemental Table S6**).

## 4. Discussion

Inflammatory changes are reported across psychiatric disorders. Here we show that specific peripheral inflammatory variables show distinct relationships with three transdiagnostic dimensions of psychopathology, namely ‘social withdrawal’, ‘anxious-depression’, and ‘compulsivity and intrusive thought’. We used a relatively homogeneous population of healthy control female participants, which helped reduce external variability and maximised our ability to detect relationships between transdiagnostic dimensions of mental health and inflammatory variables at both baseline and in response to stress. However, this approach may have limited the extent to which our findings generalise to other populations. Therefore, this work should be considered a starting point for future studies linking transdiagnostic psychopathology with inflammatory changes. Nevertheless, our data are largely consistent with previous reports in both human (Fauci and Dale, 1974; Heidt et al., 2014; Kerr, 1956; Maydych et al., 2017) and animal models (Curtin et al., 2009, 2009; Dhabhar et al., 1994; Panzenhagen and Speirs, 1953) that collectively suggest acute stress activates the innate immune system (i.e., neutrophils, monocytes) while simultaneously suppressing the adaptive immune system (i.e., lymphocytes).

Specifically, we found strong evidence for increased levels of neutrophils post-stress, the magnitude of which was positively associated with the ‘anxious-depression’ factor. At baseline, neutrophil counts were furthermore associated with ‘social withdrawal’. Neutrophilia has consistently been reported as a consequence of stress exposure in humans and in animal models (Ao et al., 2020; Heidt et al., 2014; Ishikawa et al., 2021; Meng et al., 2024), driven by rapid release of neutrophils from bone marrow stores by the sympathetic nervous system. In addition, the current study is consistent with our previous demonstration in animal models that blood neutrophils are related to anxiety- and depressive-like behaviour (Kigar et al., 2025; Lynall et al., 2021). The link between increased neutrophils post-stress and the ‘anxious-depression’ factor is particularly interesting: this factor has been reliably related to lower confidence in perceptual decision-making, indicating a metacognitive alteration in decision-making (Rouault et al., 2018). Cognitive behavioural therapy (CBT) has specifically been shown to reduce patients’ scores on this transdiagnostic factor, with reductions associated with improvements in metacognitive confidence (Fox et al., 2023). Building on this, future work could examine the capacity for psychological interventions like CBT to ameliorate stress-induced increases in neutrophils, or use stress-induced increases in neutrophils to predict CBT treatment response.

While overall blood monocyte levels did not change from baseline to post-stress, individual baseline variation in monocyte levels was related to two of our transdiagnostic dimensions: ‘social withdrawal’ scores, which were higher in individuals with higher levels of monocytes, and ‘anxious-depression’ factor scores, which were lower in individuals with higher monocyte counts. The positive relationship between monocytes and ‘social withdrawal’ is further supported by our principal component analysis of inflammatory variables, which showed a relationship between social withdrawal and immune PC2, which itself was weighted most strongly for monocytes. This relationship between monocyte levels and ‘social withdrawal’ is consistent with work in the social defeat animal model for depression (Hodes et al., 2014), which uses reduced investigation of a novel social stimulus to assess social anhedonia, finding elevated baseline monocyte levels are negatively correlated with social investigation. In humans, the ‘social withdrawal’ factor is specifically associated with biased planning during social decision-making (Hunter et al., 2022), suggesting cognitive as well as inflammatory associations with this transdiagnostic profile.

We additionally found an increase in IL-6^+^ monocytes in LPS-stimulated PBMCs from participants collected post-stress compared to baseline, which is consistent with previous reports in chronically-stressed caregivers (Miller et al., 2014). Moreover, the magnitude of change in plasma IL-6 levels from baseline to post-stress was positively related to ‘anxious-depression’. This is consistent with animal models of chronic stress-associated depression (Hodes et al., 2014; Voorhees et al., 2013). Others have shown that LPS-stimulation of blood collected from stressed participants leads to increased IL-6 release, as detected in the supernatant (Maydych et al., 2017; Miller and Chen, 2010). However, in these studies, the source of IL-6 was not directly assessed, which is important given that IL-6 can be produced by a variety of cell types (Grebenciucova and VanHaerents, 2023). Our study addresses this issue by directly interrogating the intracellular concentration of IL-6 in peripheral white blood cells using flow cytometry, enabling positive identification of monocytes as the cell type responding to psychosocial stress. Identification of the cellular source of proinflammatory IL-6 may facilitate future therapies for psychiatric disorders like depression (Osimo et al., 2020), eating disorders (Dalton et al., 2018), or PTSD (Passos et al., 2015)—all of which show elevated circulating levels of IL-6. Recent work has also highlighted a multi-protein-derived measure of IL-6 activity as a potential biomarker linked to clinical and cognitive outcomes in depression (Foley et al., 2024). Future work adapting the present methodology may clarify the contribution of monocytes and other cell types to IL-6 production across specific psychiatric disorders.

Conversely, we observed a decrease in lymphocytes, eosinophils, and IFN-γ levels when comparing baseline to post-stress blood samples, which is consistent with previous reports (Curtin et al., 2009; Fauci and Dale, 1974; Kerr, 1956; Maydych et al., 2017). IFN-γ is primarily secreted by T and NK cell lymphocytes (Schoenborn and Wilson, 2007); reduced plasma levels of IFN-γ may therefore reflect the observed decrease in overall lymphocytes. Notably, low IFN-γ has previously been associated with depressive symptoms (Daria et al., 2020), underscoring its potential relevance to stress-related pathophysiology. Eosinophils are technically part of the innate immune system, but their levels correlate with lymphocytes in a variety of disease states (Baumann et al., 2025; Carroll et al., 1997; Ownby et al., 1983; Shah et al., 2016), suggesting co-regulatory mechanisms for these cell types.

In the current study, we are unable to discern between different lymphocyte subpopulations in blood (i.e., T cells, NK cells, B cells, etc.), which may dilute any relationships with transdiagnostic dimensions of psychopathology. For example, while we did not find support for relationships between lymphocytes or IFN-γ and transdiagnostic dimensions in the current study, this may relate to the granularity of the cell subsets assayed. In previous reports of relationships between lymphocytes and stress-associated behavioural change or stress-associated disorders, shifts in lymphocyte subpopulations have been more marked than effects on the overall number of lymphocytes (Kigar et al., 2025; Lynall et al., 2021). This is underscored by our observation in the current study that plasma IL-12p70 levels are negatively related to compulsivity and intrusive thought. This relationship was observed at baseline, and when accounting for stress-induced differences in IL-12p70 levels. IL-12p70 is a cytokine produced by antigen presenting cells of the innate immune system (e.g. macrophages, dendritic cells, etc.) which polarises CD4^+^ T cells toward an IFN-γ-producing Th1 phenotype (Gee et al., 2009). Interestingly, IL-12p70 levels have previously been linked to psychiatric disorders with compulsivity as a core symptom—specifically, Tourette’s syndrome (Yeon et al., 2017) and anorexia nervosa (Keeler et al., 2021). Cognitively, the ‘compulsivity and intrusive thought’ dimension is reliably and specifically linked to a shift towards habitual, over goal-directed decision-making (Gillan et al., 2016) — a cognitive feature observed across binge eating, methamphetamine dependence, and obsessive-compulsive disorder (Voon et al., 2015) or behavioural addictions (Solly et al., 2025). Future efforts to use a deeper immunophenotyping panel to capture variation in the adaptive immune system in relation to transdiagnostic factors, ideally also capturing cognitive profiles, is warranted.

The results of this study should be considered in light of several limitations. The sample size was modest, which may have limited power to detect small effects and constrained the generalisability of findings. In addition, the sample consisted exclusively of healthy female participants, with the aim of reducing variability and allowing clearer examination of associations between transdiagnostic dimensions of mental health and inflammatory responses. However, this focus also constrains the generalisability of our findings. By focusing on healthy participants, we cannot determine whether the same relationships would emerge, or whether the direction or magnitude may differ, in individuals experiencing greater symptom severity. Moreover, as the sample was drawn from a mostly Western European population, the findings may not generalise to individuals from different cultural backgrounds.

The decision to have a female-only sample was motivated by the goal of minimising physiological confounds in a small biological study. Historically, biomedical research, both in humans and in animal models, has disproportionately focused on males, partly due to concerns that ovarian hormone variation would increase variability (Becker et al., 2016; Shansky and Murphy, 2021). This bias has limited the representation of female physiology in medical research. Despite this, men also show diurnal variation in testosterone and cortisol levels (Cooke et al., 1993; de la Torre et al., 1981; Diver et al., 2003), which can similarly confound inflammatory levels. Restricting the sample to a single sex therefore improved internal consistency, though future work should include both male and female participants, individuals with greater symptom severity, as well as more diverse cultural backgrounds, to determine whether similar associations are observed and to promote a more equitable understanding of the biological pathways linking mental health and inflammation.

## 5. Conclusions

In summary, our findings provide novel evidence that distinct peripheral immune signatures are differentially associated with transdiagnostic dimensions of psychopathology, both at baseline and following acute stress. By moving beyond traditional categories and employing a dimensional framework, we disentangled the shared and unique correlates of ‘social withdrawal’, ‘anxious-depression’, and ‘compulsivity and intrusive thought’, revealing patterns that may be obscured in categorical case-control study designs. A transdiagnostic approach offers clear advantages for the field: it aligns more closely with underlying biological processes and supports the development of precision therapies that can be aimed at transdiagnostic symptoms rather than heterogeneous diagnostic categories. The use of transdiagnostic frameworks will be essential to uncovering the complex immunological underpinnings of mental health problems and for guiding the development of more targeted and effective biologically informed interventions.

## Author Contributions

Author contributions are as follows: conceptualisation and methodology: AJS, SLK, MEL, TD, CLN; investigation, AJS, SLK, QD; formal analysis, AJS, SLK, QD, CLN; project administration, KI, MK, TD; writing - original draft, AJS, SLK, CLN; writing - review and editing, AJS, SLK, QD, MEL, KI, MK, CH, TD, CLN; supervision - CH, TD, CLN; funding acquisition, CLN.

## Declaration of Competing Interest

KI is receiving a stipend from Elsevier for editorial work.

## Acknowledgements

Cytokine assays were performed by the NIHR Cambridge BRC Core Biochemical Assay Laboratory.

This research was supported by the NIHR Cambridge Biomedical Research Centre Mental Health theme (BRC-1215-20014), the Cambridge NIHR BRC Cell Phenotyping Hub, the AXA Research Fund (G102329, awarded to CLN), and the Medical Research Council (MC_UU_00030/12, awarded to CLN). CLN is funded by a Wellcome Career Development Award (226490/Z/22/Z). SLK is funded by the NIHR Cambridge Biomedical Research Centre (NIHR203312 & BRC-1215-20014). MEL is supported by the Addenbrooke’s Charitable Trust, the Medical Research Council (MR/S006257/1), and the Cambridge BRC. KI’s research is supported by the National Institute of Health and Care Research (NIHR) Applied Research Collaboration (ARC) Wessex. The views expressed are those of the author(s) and not necessarily those of the NHS, NIHR or the Department of Health and Social Care (DHSC) UK.

## Code availability

The data and code used to run the analyses and create the plots in this paper are shared openly on GitHub (https://github.com/aliciasmith1/transdiagnostic-inflammatory-profiles).

