## Supplemental Materials for "Inflammatory profiles of transdiagnostic symptom dimensions in healthy females"

**Supplemental Materials**
**Inflammatory profiles of transdiagnostic dimensions in healthy females**

Alicia J. Smith*, Stacey L. Kigar*, Quentin Dercon, Mary-Ellen Lynall, Konstantinos Ioannidis, Muzaffer Kaser, Caitlin Hitchcock, Tim Dalgleish & Camilla L. Nord

*Joint first author

**Supplemental Table S1.** Imprecision data showing cytokine mean (pg/ml) and inter-assay coefficient variation (CV) values derived from controls run across dozens of MesoScale Discovery Human 10-plex ProInflammatory Cytokine panel assays.

| Analyte | Mean | %CV | Mean | %CV | Mean | %CV |
|---|---|---|---|---|---|---|
| **IFN - ɣ** | 22.9 | 19.91 | 86.8 | 15.38 | 327 | 14.39 |
| **IL - 1β** | 7.8 | 9.79 | 30.9 | 7.42 | 124 | 7.21 |
| **IL – 2** | 22.5 | 9.37 | 89.5 | 7.34 | 348 | 7.18 |
| **IL – 4** | 3.7 | 14.33 | 14.5 | 11.33 | 55.2 | 12.93 |
| **IL – 6** | 10.4 | 10.69 | 38.5 | 7.85 | 155 | 8.94 |
| **IL – 8** | 8.1 | 7.72 | 31.4 | 6.20 | 124 | 5.90 |
| **IL – 10** | 5.1 | 10.78 | 20.1 | 9.69 | 78.8 | 8.04 |
| **IL – 12p70** | 7.1 | 14.97 | 27.9 | 11.96 | 109 | 12.25 |
| **IL – 13** | 5.6 | 13.62 | 19.6 | 8.43 | 87 | 8.30 |
| **TNF - α** | 3.9 | 13.07 | 15.0 | 11.01 | 60.6 | 8.86 |

**Supplemental Table S2.** Antibodies, blocking reagent, and viability marker used in this study.

| Antibody | Vendor | Catalogue |
|---|---|---|
| **CD56-FITC** | Invitrogen | 11-0566-42 |
| **CD123-PerCP/Cy5.5** | Invitrogen | 45-1239-42 |
| **CD11c-PE/Vio770** | Miltenyi | 130-113-581 |
| **CD16-APC** | Invitrogen | 17-0168-42 |
| **CD20-APC/Cy7** | Invitrogen | 47-0209-42 |
| **CD19-APC/Cy7** | Invitrogen | 47-0199-42 |
| **HLA-DR-e450** | Invitrogen | 48-9952-42 |
| **CD14-BV605** | BioLegend | 301834 |
| **CD3-BV650** | BioLegend | 317324 |
| **Live/Dead Aqua** | Thermo Fisher | L34957 |
| **Human Fc block** | Miltenyi | 130-059-901 |

**Supplemental Figure S1.** Full gating strategy used to identify 6 unique immune cell types. After 4h of LPS stimulation, PBMCs were stained for extracellular cell lineage markers. Cells were then fixed, permeabilized, and stained for intracellular IL6 (shown in **Figure 2**). Flow cytometry data were acquired and gated to identify the following unique cell populations: monocytes (CD14$^{+}$), B cells (CD19$^{+}$/CD20$^{+}$), T cells (CD3$^{+}$), NKT cells (CD3$^{+}$; CD56$^{+}$), NK cells (CD3$^{-}$; CD56$^{+}$), and dendritic cells (CD11c$^{+}$; HLA-DR$^{+}$).

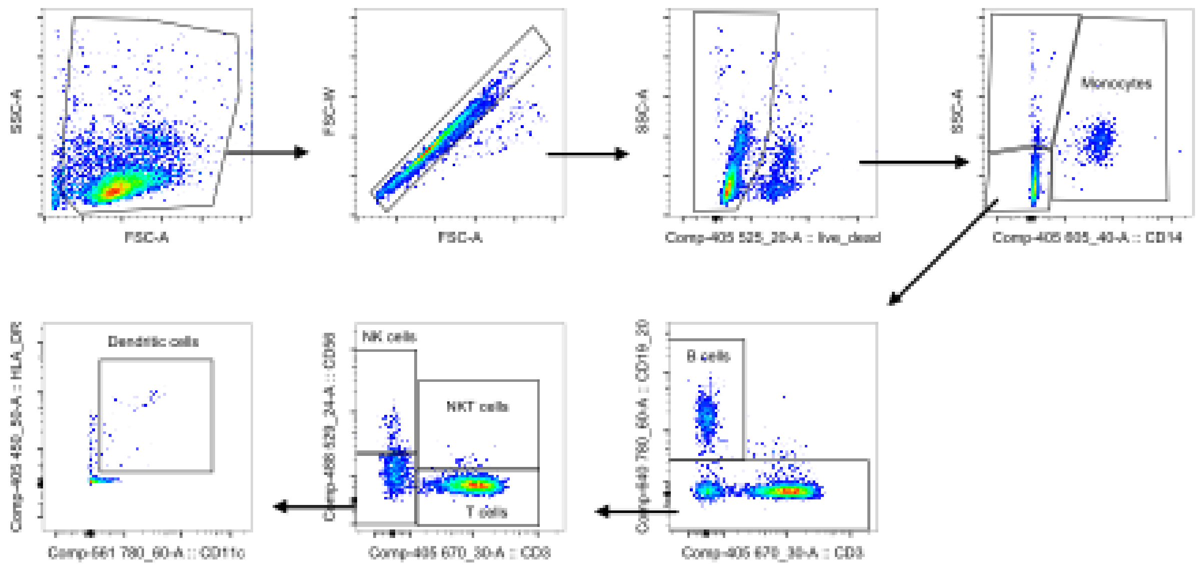


**Supplemental Table S3.** Estimates for the Bayesian regression model predicting baseline measurements of white blood cell counts, plasma cytokines and intracellular IL-6 staining from the three transdiagnostic dimensions.

| Target variable | Predictor | Estimate (+/- HDI) | Bayes Factor |
|---|---|---|---|
| **Neutrophils** | Social Withdrawal | 0.69 (0.06, 1.32) | 2.92 |
| **Neutrophils** | Anxious-Depression | -0.1 (-0.61, 0.39) | 0.27 |
| **Neutrophils** | Compulsivity & Intrusive Thought | -0.04 (-0.55, 0.46) | 0.26 |
| **Monocytes** | Social Withdrawal | 0.89 (0.29, 1.46) | 19.08 |
| **Monocytes** | Anxious-Depression | -0.5 (-0.96, -0.02) | 1.81 |
| **Monocytes** | Compulsivity & Intrusive Thought | 0.11 (-0.35, 0.59) | 0.27 |
| **Lymphocytes** | Social Withdrawal | 0.14 (-0.57, 0.86) | 0.39 |
| **Lymphocytes** | Anxious-Depression | 0.03 (-0.56, 0.59) | 0.3 |
| **Lymphocytes** | Compulsivity & Intrusive Thought | -0.15 (-0.73, 0.43) | 0.34 |
| **Eosinophils** | Social Withdrawal | -0.14 (-0.75, 0.47) | 0.34 |
| **Eosinophils** | Anxious-Depression | 0.03 (-0.46, 0.52) | 0.25 |
| **Eosinophils** | Compulsivity & Intrusive Thought | -0.23 (-0.72, 0.26) | 0.39 |
| **Basophils** | Social Withdrawal | 0.2 (-0.58, 1.0) | 0.47 |
| **Basophils** | Anxious-Depression | -0.25 (-0.89, 0.38) | 0.45 |

| | | | |
|---|---|---|---|
| **Basophils** | Compulsivity & Intrusive Thought | 0.05 (-0.61, 0.67) | 0.33 |
| **IL-6** | Social Withdrawal | -0.26 (-0.7, 0.16) | 0.45 |
| **IL-6** | Anxious-Depression | 0.25 (-0.14, 0.65) | 0.43 |
| **IL-6** | Compulsivity & Intrusive Thought | -0.2 (-0.59, 0.18) | 0.33 |
| **IL-8** | Social Withdrawal | 0.41 (-0.1, 0.93) | 0.92 |
| **IL-8** | Anxious-Depression | -0.02 (-0.5, 0.46) | 0.25 |
| **IL-8** | Compulsivity & Intrusive Thought | -0.06 (-0.53, 0.4) | 0.24 |
| **IL-10** | Social Withdrawal | -0.08 (-0.5, 0.31) | 0.23 |
| **IL-10** | Anxious-Depression | -0.01 (-0.38, 0.38) | 0.19 |
| **IL-10** | Compulsivity & Intrusive Thought | -0.22 (-0.59, 0.14) | 0.38 |
| **TNF-a** | Social Withdrawal | -0.24 (-0.65, 0.16) | 0.4 |
| **TNF-a** | Anxious-Depression | 0.04 (-0.34, 0.41) | 0.2 |
| **TNF-a** | Compulsivity & Intrusive Thought | 0.19 (-0.16, 0.56) | 0.32 |
| **INF-y** | Social Withdrawal | -0.26 (-0.61, 0.09) | 0.49 |
| **INF-y** | Anxious-Depression | -0.03 (-0.36, 0.3) | 0.17 |
| **INF-y** | Compulsivity & Intrusive Thought | -0.19 (-0.5, 0.12) | 0.34 |
| **IL-12p70** | Social Withdrawal | 0.12 (-0.24, 0.51) | 0.23 |
| **IL-12p70** | Anxious-Depression | 0.21 (-0.14, 0.54) | 0.35 |
| **IL-12p70** | Compulsivity & Intrusive Thought | -0.36 (-0.69, -0.02) | 1.39 |
| **IL6 Monocytes** | Social Withdrawal | 0.23 (-0.54, 1.04) | 0.46 |
| **IL6 Monocytes** | Anxious-Depression | -0.33 (-1.14, 0.47) | 0.59 |
| **IL6 Monocytes** | Compulsivity & Intrusive Thought | 0.52 (-0.2, 1.25) | 1.03 |
| **IL6 B Cells** | Social Withdrawal | 0.15 (-0.64, 0.93) | 0.43 |
| **IL6 B Cells** | Anxious-Depression | -0.53 (-1.36, 0.27) | 0.99 |
| **IL6 B Cells** | Compulsivity & Intrusive Thought | 0.7 (-0.04, 1.42) | 2.11 |

**Supplemental Table S4.** Estimates for the Bayesian regression model predicting differences in white blood cell counts, plasma cytokines and intracellular IL-6 staining between baseline and post-stress from the three transdiagnostic dimensions.

| Target variable | Predictor | Estimate (+/- HDI) | Bayes Factor |
|---|---|---|---|
| **Neutrophils** | Social Withdrawal | -0.12 (-0.86, 0.62) | 0.39 |
| **Neutrophils** | Anxious-Depression | 1.14 (0.18, 2.14) | 5.66 |
| **Neutrophils** | Compulsivity & Intrusive Thought | -0.22 (-0.99, 0.6) | 0.48 |

| | | | |
|---|---|---|---|
| **Monocytes** | Social Withdrawal | 0.35 (-0.64, 1.3) | 0.65 |
| **Monocytes** | Anxious-Depression | -0.14 (-1.45, 1.17) | 0.69 |
| **Monocytes** | Compulsivity & Intrusive Thought | 0.05 (-1.03, 1.1) | 0.54 |
| **Lymphocytes** | Social Withdrawal | -0.06 (-1.01, 0.86) | 0.49 |
| **Lymphocytes** | Anxious-Depression | -0.03 (-1.28, 1.21) | 0.64 |
| **Lymphocytes** | Compulsivity & Intrusive Thought | 0.16 (-0.83, 1.17) | 0.54 |
| **Eosinophils** | Social Withdrawal | -0.18 (-0.86, 0.56) | 0.42 |
| **Eosinophils** | Anxious-Depression | -0.25 (-1.22, 0.68) | 0.56 |
| **Eosinophils** | Compulsivity & Intrusive Thought | 0.37 (-0.39, 1.14) | 0.65 |
| **Basophils** | Social Withdrawal | 0.32 (-0.95, 1.6) | 0.74 |
| **Basophils** | Anxious-Depression | 0.02 (-1.65, 1.73) | 0.84 |
| **Basophils** | Compulsivity & Intrusive Thought | -0.35 (-1.73, 1.0) | 0.81 |
| **IL-6** | Social Withdrawal | -0.6 (-1.3, 0.11) | 1.48 |
| **IL-6** | Anxious-Depression | 1.09 (0.03, 2.17) | 3.59 |
| **IL-6** | Compulsivity & Intrusive Thought | -0.53 (-1.39, 0.29) | 0.93 |
| **IL-8** | Social Withdrawal | -0.16 (-0.92, 0.6) | 0.43 |
| **IL-8** | Anxious-Depression | -0.29 (-1.46, 0.89) | 0.68 |
| **IL-8** | Compulsivity & Intrusive Thought | 0.19 (-0.74, 1.12) | 0.53 |
| **IL-10** | Social Withdrawal | 0.43 (-0.24, 1.12) | 0.79 |
| **IL-10** | Anxious-Depression | -0.04 (-1.08, 1.02) | 0.55 |
| **IL-10** | Compulsivity & Intrusive Thought | -0.46 (-1.28, 0.36) | 0.79 |
| **TNF-a** | Social Withdrawal | 0.25 (-0.36, 0.85) | 0.44 |
| **TNF-a** | Anxious-Depression | -0.03 (-0.94, 0.91) | 0.49 |
| **TNF-a** | Compulsivity & Intrusive Thought | -0.48 (-1.12, 0.26) | 0.86 |
| **INF-y** | Social Withdrawal | 0.2 (-0.31, 0.72) | 0.36 |
| **INF-y** | Anxious-Depression | 0.09 (-0.7, 0.88) | 0.42 |
| **INF-y** | Compulsivity & Intrusive Thought | 0.15 (-0.46, 0.77) | 0.36 |
| **IL-12p70** | Social Withdrawal | 0.3 (-0.2, 0.8) | 0.51 |
| **IL-12p70** | Anxious-Depression | 0.12 (-0.62, 0.9) | 0.41 |
| **IL-12p70** | Compulsivity & Intrusive Thought | -0.69 (-1.29, -0.09) | 3.49 |
| **IL6 Monocytes** | Social Withdrawal | -0.18 (-1.08, 0.78) | 0.51 |
| **IL6 Monocytes** | Anxious-Depression | 0.12 (-1.48, 1.62) | 0.81 |
| **IL6 Monocytes** | Compulsivity & Intrusive Thought | -0.24 (-1.38, 0.89) | 0.63 |
| **IL6 B Cells** | Social Withdrawal | -0.27 (-1.23, 0.7) | 0.56 |
| **IL6 B Cells** | Anxious-Depression | 0.92 (-0.68, 2.52) | 1.58 |
| **IL6 B Cells** | Compulsivity & Intrusive Thought | -0.15 (-1.13, 1.01) | 0.62 |

**Supplemental Figure S2.** A bar plot displaying the variance explained by each principal component derived from inflammatory measurements at (A) baseline, and (B) in terms of pre- to post-stress differences.

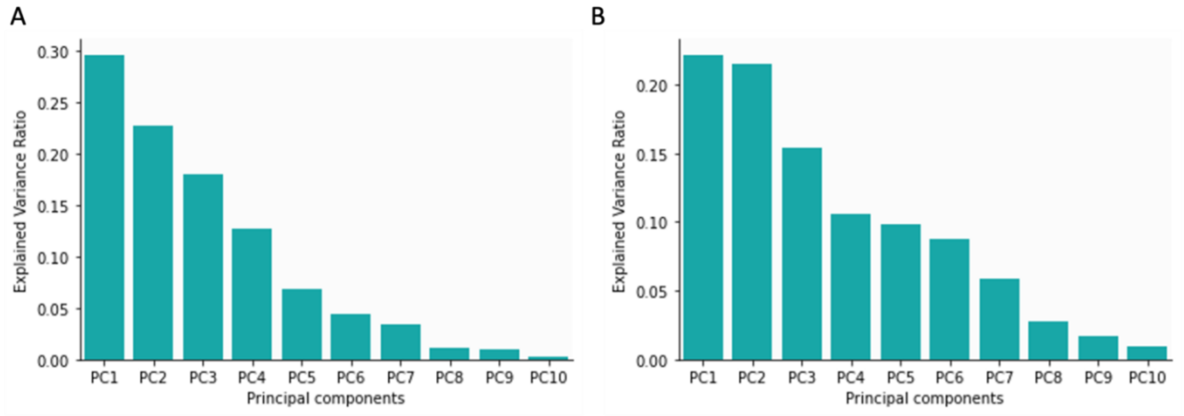


**Supplemental Table S5.** Estimates for the Bayesian regression model for which the three transdiagnostic factors predict scores on PC1, PC2 and PC3 derived from measurements of inflammatory variables at baseline.

| Target variable | Predictor | Estimate (+/- HDI) | Bayes Factor |
|---|---|---|---|
| **PC1** | Social Withdrawal | 0.28 (-0.19, 0.74) | 0.51 |
| **PC1** | Anxious-Depression | -0.17 (-0.55, 0.22) | 0.31 |
| **PC1** | Compulsivity & Intrusive Thought | 0.15 (0.16, 0.45) | 0.26 |
| **PC2** | Social Withdrawal | -0.55 (-1.14. -0.01) | 1.88 |
| **PC2** | Anxious-Depression | -0.12 (-0.61, 0.34) | 0.27 |
| **PC2** | Compulsivity & Intrusive Thought | 0.27 (-0.09, 0.66) | 0.56 |
| **PC3** | Social Withdrawal | -0.07 (-0.72, 0.6) | 0.33 |
| **PC3** | Anxious-Depression | -0.2 (-0.74, 0.35) | 0.37 |
| **PC3** | Compulsivity & Intrusive Thought | 0.08 (-0.33, 0.52) | 0.23 |

**Supplemental Table S6.** Estimates for the Bayesian regression model for which the three transdiagnostic factors predict scores on PC1, PC2 and PC3 derived from measurements of inflammatory variables in terms of their pre- to post-stress differences.

| Target variable | Predictor | Estimate (+/- HDI) | Bayes Factor |
|---|---|---|---|
| **PC1** | Social Withdrawal | -0.5 (-1.07, 0.05) | 1.35 |
| **PC1** | Anxious-Depression | 0.4 (-0.06, 0.86) | 5.02 |
| **PC1** | Compulsivity & Intrusive Thought | -0.03 (-0.4, 0.34) | 3.7 |
| **PC2** | Social Withdrawal | 0.04 (-0.45, 0.56) | 0.95 |
| **PC2** | Anxious-Depression | 0.35 (-0.06, 0.77) | 3.9 |

| | | | |
|---|---|---|---|
| **PC2** | Compulsivity & Intrusive Thought | -0.28 (-0.61, 0.04) | 2.74 |
| **PC3** | Social Withdrawal | -0.09 (-0.66, 0.46) | 1.45 |
| **PC3** | Anxious-Depression | 0.26 (-0.21, 0.71) | 6.32 |
| **PC3** | Compulsivity & Intrusive Thought | -0.17 (-0.53, 0.2) | 4.63 |
